# In vitro detection of canine anti-human antibodies following intratumoral injection of the hu14.18-IL2 immunocytokine in spontaneous canine melanoma

**DOI:** 10.1101/2025.03.21.644578

**Authors:** Andrew T. Kosharek, Cindy L. Zuleger, William S. Glass, Jens Eickhoff, Paul M. Sondel, David M. Vail, Mark R. Albertini

## Abstract

**Background:** Canine and human malignant melanoma are naturally occurring cancers with many similarities, making the dog an important parallel patient population to study both diseases. However, development of canine anti-human antibodies (CAHA) needs to be considered when evaluating humanized biotherapeutics in dogs.

**Objectives:** Characterize CAHA in sera from dogs with spontaneous melanoma receiving radiotherapy and intratumoral immunocytokine (IT-IC) with humanized 14.18-IL2.

**Methods:** Serum samples were obtained pre-treatment and at several post-treatment times from 12 dogs with locally advanced or metastatic melanoma treated with radiotherapy to the primary site and regional lymph nodes (when clinically involved) followed by IT-IC of humanized 14.18-IL2. Two CAHA assays were developed. A sandwich enzyme-linked immunosorbent assay (ELISA) was developed to detect antibodies against the humanized IgG component of hu14.18-IL2. A flow cytometry assay was developed to determine the ability of CAHA to inhibit binding of a mouse anti-GD2 monoclonal antibody to its target.

**Results:** Post-treatment sera from 7 of 12 dogs developed CAHA levels over pre-treatment that were identified by ELISA as significant increases at Day 30 and/or Day 60. Day 10, Day 30, and Day 60 post-treatment sera from 10 of 12 dogs significantly inhibited the binding of anti-GD2 monoclonal antibody to its target compared to pre-treatment. Significant binding inhibition was also detected in 2 of 12 dogs after local RT but before IT-IC (Day 1). Normal canine sera did not mediate binding inhibition.

**Conclusions:** This study advances CAHA detection strategies and reports the kinetics of CAHA following IT-IC in dogs with spontaneous melanoma.

## Introduction

Companion (pet) dogs with spontaneous tumors provide unique and clinically relevant insights into human tumor biology, therapy, and drug development [1, 2]. Dogs develop spontaneous tumors, similar to humans [3]. This contrasts with many laboratory mouse models where tumors are experimentally induced [4]. Moreover, the molecular and genetic characteristics of many canine cancers resemble those of human cancers [5]. While no model perfectly recapitulates human cancer, the companion dog melanoma patient population holds a distinctive position in oncology research and has received attention from the scientific community as being somewhat parallel to human melanoma patients, particularly for evaluation of immunotherapy studies [6–10]. In that regard, development of antidrug antibodies (ADAs) against immunotherapeutic monoclonal antibodies (mAbs) is well documented in human patients, and the presence of ADAs can impact the safety and efficacy of immunotherapy [11–15]. Moreover, the immune response is highly species-specific and relatively few immunotherapeutic mAbs are caninized, i.e., canine speciated [16]. Therefore, it will be critical to monitor ADAs in the setting of canine immunotherapy, particularly when the immunotherapeutic is a mAb of a species ‘foreign’ to the dog (e.g., a humanized mAb).

The humanized mAb Hu14.18-IL2 and the chimeric ch14.18-IL2, both GD2-targeting mAbs linked to IL2 and known as immunocytokines (IC), have been studied *in vitro* and in preclinical mouse models with GD2-expressing tumors [17–20]. Clinical trials evaluating intravenous (IV) delivery of hu14.18-IL2 IC in patients with melanoma or neuroblastoma have demonstrated immune activation and had manageable safety profiles [21–25]. Although hu14.18-IL2 is a humanized mAb, some patients still developed antibodies to the therapeutic IC [22, 26]. In these patients, antibodies were detected to both the idiotype, i.e., the antigen binding end, and the Fc-IL2 end of the IC [22, 26]. A substantial fraction of patients develop a detectible anti-IC antibody response, and some patients develop a strong anti-IC response and functional, in vivo “neutralization” of detectible IC. Namely, following detection of a strong anti-IC response, blood obtained immediately after an IC infusion does not contain the anticipated level of detectible IC, based on the IC level in the blood obtained from samples collected prior to development of the anti-IC response [22, 26]. A neutralizing anti-human antibody (HAHA) or anti-chimeric antibody (HACA) has been reported for some patients treated with humanized or chimeric hu14.18 mAb [27, 28].

The immunogenicity of the humanized hu14.18-IL2 is potentially a greater concern when administering this molecule to a “foreign” species, e.g., to a mouse or dog, as ADAs could interfere with the pharmacokinetics of this therapy, potentially reducing its efficacy [29]. When huKS1/4-IL2, a humanized antibody directed against the human epithelial cell adhesion molecule, was administered intravenously to mice with adenocarcinoma, the mice mounted significant ADAs in the form of mouse anti-human antibodies (MAHAs) resulting in rapid clearance of additional doses of the therapeutic [18]. Delivering hu14.18-IL2 intratumorally in mouse models also induced a MAHA response, similar to what was seen following IV-IC therapy [30]. In addition, ADAs in dogs have been observed in other studies, including one report following subcutaneous administration of a therapeutic IC [31–33]. However, the development of canine anti-human antibodies (CAHAs) following intratumoral immunotherapy with an IC has not been reported.

Here we present CAHA assay development and CAHA monitoring of companion dogs with spontaneous melanoma enrolled in a clinical trial of local RT plus intratumoral hu14.18-IL2. An in-house ELISA was developed and used to monitor CAHA against the IgG portion of the IC. Previous studies have demonstrated the ability of ELISA to detect ADAs against antibody-based biopharmaceuticals [26, 34–37]. Our ELISA is customizable, cost-effective, and uses common laboratory reagents and equipment. We also examined if CAHA interferes with the ability of the GD2-binding mAb part of the IC to recognize GD2 in a cell-based flow cytometry assay. Details about safety, clinical activity, and tumor biopsy analyses from this trial, as well as another trial, are separately reported [38, 39].

## Material and Methods

### Trial eligibility and enrollment

Client-owned companion (pet) dogs weighing ≥5 kg with a readily palpable and accessible oral malignant melanoma (histologically confirmed) of at least 2.0 cm (longest diameter) were included in the study. Both treatment naïve dogs and dogs with surgically recurrent tumors were allowed, provided the recurrence met the minimum size criteria. All dogs were required to have a pretreatment constitutional clinical sign status of 0 or 1 (normal or mild lethargy over baseline; diminished activity from pre-disease level but able to function as an acceptable pet) according to VCOG-CTCAE v2 criteria at study entry [40]. Dogs determined to have any significant co-morbid illness were excluded.

### Study design

Client-owned companion dogs with locally advanced or metastatic melanoma were randomized (6 dogs per Arm) to receive either a single 8 Gy fraction (Arm A) or three 8 Gy fractions delivered over 1 week (Arm B) to the primary site and regional lymph nodes (when clinically involved) with the single or last fraction 5 days prior to IT-IC at 12 mg/m^2^ on 3 consecutive days. Prior to treatment, a baseline physical examination with tumor measurements was performed, blood was collected for clinical assessments, and blood was collected for banking of serum, plasma, and peripheral blood mononuclear cells (PBMC) for subsequent *in vitro* analysis. External beam radiation therapy (RT) was delivered by image-guided intensity modulated radiation using helical tomotherapy (TomoTherapy HiArt Treatment System, Accuray Inc., Sunnyvale, CA, USA). The hu14.18-IL2 was provided as lyophilized vials (4 mg/vial; each ml prior to lyophilization contained 4 mg/ml hu14.18-IL2, 2% sucrose, 80mM L-arginine, 10mM citric acid, 0.2% polysorbate 20, pH 5.5) and was reconstituted with either 0.5 ml or 0.25 ml 0.9% (w/v) NaCl for injection at a concentration of either 8 mg/ml or 16 mg/ml. The total volume was administered using multiple needle re-directions into the target lesion within and surrounding the lesion. Injections were performed on 3 consecutive days. PBMC were obtained at pre-treatment and various times post-treatment. Etomidate (1-2 mg/kg, IV) and/or Propofol (4-5 mg/kg IV) were used to induce anesthesia in dogs receiving RT. Isoflurane (1.2-1.3%, inhalation) and/or Sevoflurane (2.4%, inhalation) were used to maintain anesthesia and the gas percentages were adjusted depending on the canine patient’s clinical response to anesthesia under guidance of the clinician. The duration of anesthesia was approximately 30 minutes. Butorphanol (0.2-0.4 mg/kg IM or IV) and/or midazolam (0.2 mg/kg IV) were used to sedate the dogs during IT-IC administration, the duration of which was approximately 45 minutes. Pulse, respiratory and heart rates were monitored throughout anesthesia and at least every 15 minutes until the dog was ambulatory.

Lidocaine was administered at the site of the IT-IC administration for pain (0.5-0.75 ml 2% Lidocaine, intradermal). Following IT-IC administration, pain was alleviated with analgesics for 3 days or longer if the clinician or owner perceived that the dog was experiencing continued discomfort. Analgesics included either Carprofen (2.2 mg/kg, orally, 2x daily), Deracoxib (3-4 mg/kg, orally, 1x daily), Meloxicam (0.1-0.2 mg/kg, orally, 1x daily), and/or Tramadol (1-5 mg/kg, orally, every 8 hours). Physical examination and canine patient history were assessed at each clinic visit. Dogs were monitored by the dog’s owner and clinician for animal health and behavior, as well as the development of adverse events (AE). AEs were graded and attributed according to VCOG-CTCAE v2 [40]. The attending clinician prescribed appropriate treatment and/or supportive care if an AE was observed. All efforts were made to minimize suffering. The study endpoint was the time the dog developed progressive disease as determined by physical examination and/or thoracic radiographs. Owners could remove a dog from the study for any reason, or the attending clinician could remove a dog from the study if it was determined there were severe AEs not manageable with supportive care.

Euthanasia was not a study endpoint nor was it part of this study and was only performed at the request of a dog’s owner. If requested, the American Veterinary Medical Association guidelines were followed [41]. The treatment schema and blood collection times are shown in Fig 1.

**Fig 1.**
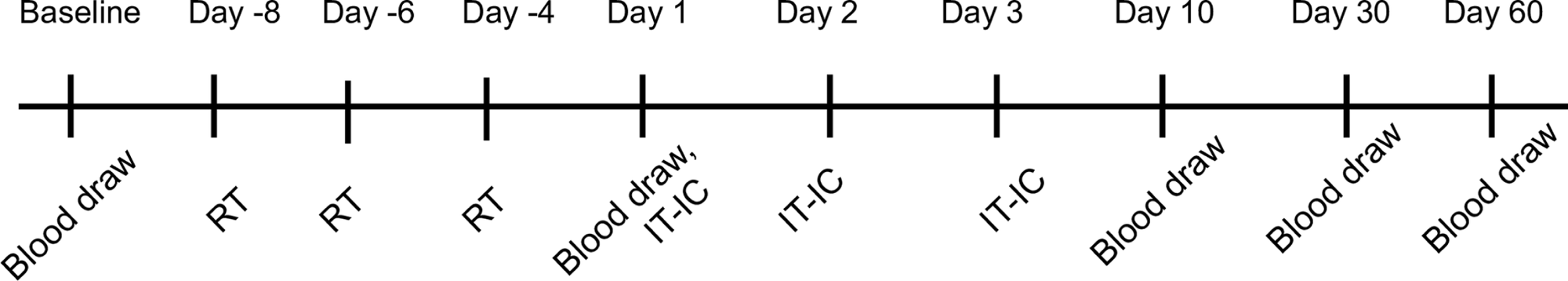
Treatment Schema. All dogs receive radiation therapy (RT) to the local site either in 1x 8 Gy (Arm A) or 3x 8 Gy fractions over 1 week (Arm B). RT was delivered on Day-4 (Arm A) or on Days-8,-6-4 (Arm B). Intratumoral (IT) injection of IC is administered (IT-IC) on three consecutive days starting five days after completion of RT. All days were allowed ± 1-2 days leeway for flexible scheduling around holidays, subject vacations, and clinic scheduling.

### IACUC approvals

The authors confirm that the appropriate ethical review committee approval has been received. All procedures and treatments performed on client-owned companion dogs were approved by the Institutional Animal Care and Use Committees of the University of Wisconsin-Madison School of Veterinary Medicine (Approval V006037) and by the Animal Care Committee at the William S. Middleton Memorial Veterans Hospital. Written informed consent was obtained from all companion dog caregivers prior to entry into this trial. Protocol treatments were administered at the University of Wisconsin Veterinary Care (UWVC) hospital.

### Canine Anti-Human ELISA

A standard protocol was adapted to develop an ELISA to detect canine antibodies against human IgG [42]. Nunc MaxiSorp™ Flat-Bottom 96-well plates (Thermo Scientific Inc., USA) were coated with ChromPure Human IgG (Jackson ImmunoResearch Laboratories, Inc., USA) [10mg/mL, 50μL/well in ELISA Coating Buffer (Bio-Rad Laboratories, Inc., USA)], which served as the “capture” agent, and incubated overnight at 4°C. The coating solution was removed, and the wells washed three times (3x) with distilled water (DI). The plate was blocked with Neptune™ Block (ImmunoChemistry Technologies, USA) (200μL/well) for 1 hour at 25°C followed by washing as above. Canine serum was diluted 1:20 in PBS and added to wells in quadruplicate at 50μL/well and incubated for 1 hour at 25°C. A murine mAb against human IgG (ab436, Abcam, USA) was used as a positive control, as we are not aware of any commercially available canine anti-human IgG to serve as a positive control. The mAb was diluted 1:1,000 in PBS and added to wells in quadruplicate at 50μL/well and incubated for 1 hour at 25°C. PBS was added to wells as a negative control. Wells were washed as above. Rabbit anti-mouse IgG (H+L) antibody, affinity purified against human serum proteins conjugated to horseradish peroxidase (HRP) (# 315-035-045, Jackson ImmunoResearch Laboratories Inc., USA) was the secondary (i.e.: “detection”) antibody, was diluted 1:40,000 in PBS, added to wells at 75μL/well, and incubated for 30 minutes (min.) at 25°C. Wells were washed as above. The plate was developed with1-Step™ Ultra TMB-ELISA solution (Thermo Scientific Inc., USA) (75μL/well) and incubated for 15 min. at 25°C. The reaction was stopped by addition of TMB stop solution (Sera Care, USA) (75μL/well), and the absorbance at 450nm read on a SpectraMax M3 Micro-Plate reader.

### Cell culture

The GD2+ M21 human melanoma cell line was cultured in RPMI-1640 supplemented with 10% fetal bovine serum (GeminiBio, BenchMark™), 25 mM HEPES, 2 mM L-glutamine, 1 mM sodium pyruvate, and 1X non-essential amino acids (Mediatech). M21 was obtained as a gift from Dr. Ralph Reisfeld (Scripps Research Institute, La Jolla, CA, USA).

### Detection of GD2 binding inhibition by CAHA using flow cytometry

Mouse anti-GD2 mAb 14G2a-PE (BioLegend, #357303) (100 μl) was added to 2.5 μl canine sera from this study (Baseline, Day 1, Day 10, Day 30, Day 60) or to serum from healthy dogs (2.5 μl or 20 μl) (Antibodies Inc, #54-490-0005) and adjusted to a total volume of 120 μl using 1% FBS/PBS flow cytometry wash buffer (WB). A sample without canine sera was used as a staining control. The light-protected mixture was incubated for 30 min. at 4°C. The serum-14G2a mixture was added to GD2+ M21 cells aliquoted to flow cytometry tubes and incubated for 30 min.at 4°C, followed by 2x washes with WB. Cells were fixed with Cytofix Fixation Buffer (BD, #554655) for 15 min. at 25°C, washed 1x with WB, resuspended in WB, and held at 4°C until analysis. Data were acquired on a BD LSR Fortessa, single cells were gated, and the Median Fluorescence Intensity (MFI) was determined with FlowJo^TM^ Software (BD Life Sciences). Binding inhibition was calculated as follows where X equals Day 1, Day 10, Day 30, or Day 60:

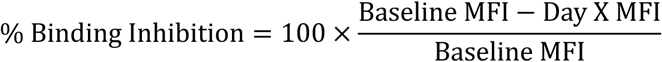

## Statistical Analysis and Considerations

Animal specific kinetic levels of CAHA following IT-IC were summarized in terms of means and standard errors. Profile plots were generated to examine changes over time. Changes from baseline to Day 10, 30, and 60 were evaluated using a generalized linear model. Model assumptions were validated by examining residual and normal probabilities plots. Dunnett’s test was used to control the type I error (<0.05) when conducting multiple comparisons for changes from baseline to Day 10, 30, and 60. All reported P values are two-sided and P<0.05 was used to define statistical significance. Statistical analyses were conducted using SAS software (SAS Institute, Cary NC), version 9.4. Plots were generated using R software version 4.2.1 with graphical packages [43–45].

## Results

### Companion dog characteristics

Twelve dogs met eligibility criteria for the trial and were enrolled and randomized into one of the two treatment groups from May 2020 to December 2021 [38]. Companion dog breed, tumor characteristics, and randomization results are summarized in Table 1.

**Table 1.**
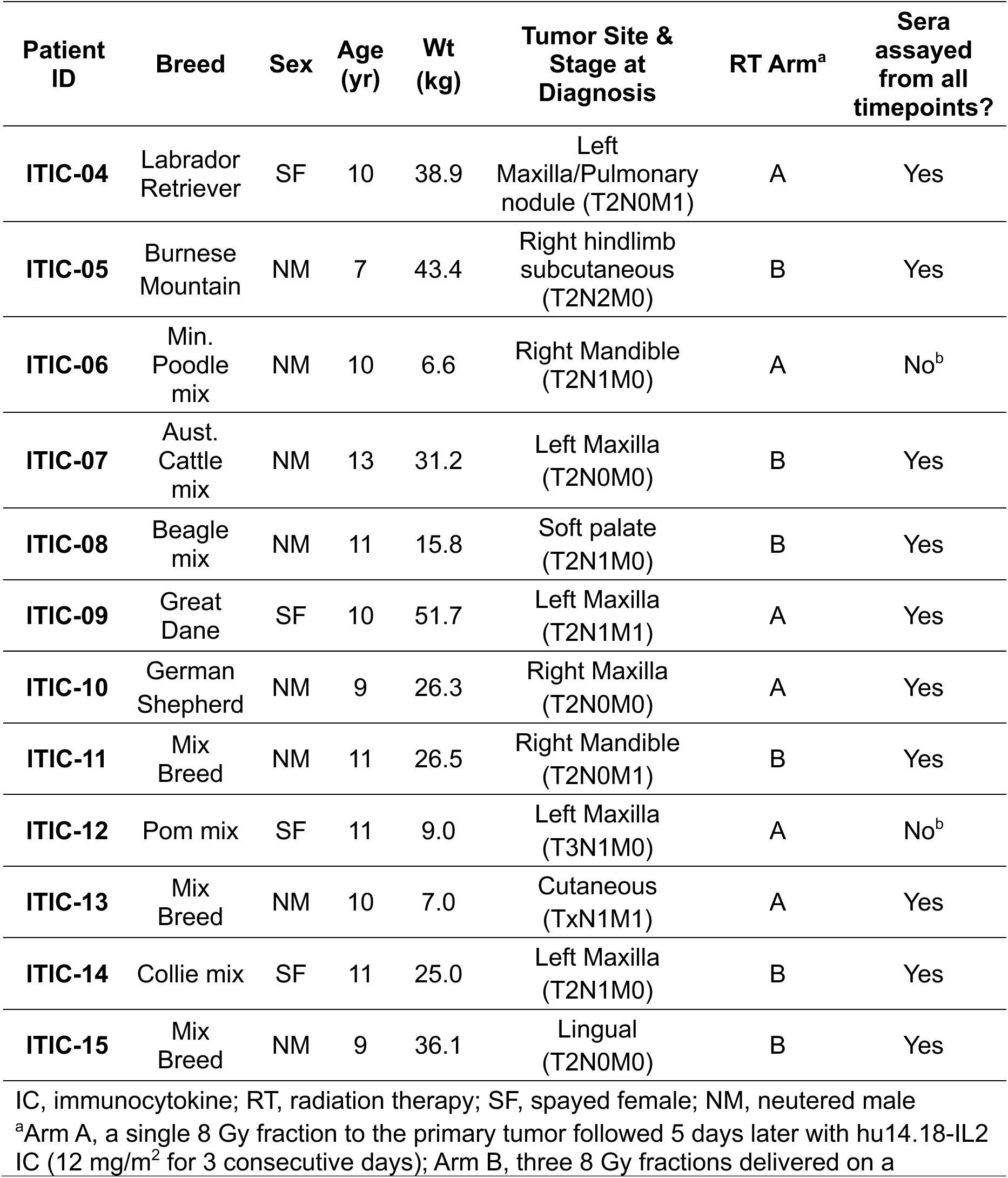
Characteristics and status of dogs receiving 12 mg/m^2^ IC and randomized to RT Arm A or RT Arm B.

| Patient ID | Breed | Sex | Age (yr) | Wt (kg) | Tumor Site & Stage at Diagnosis | RT Arm<sup>a</sup> | Sera assayed from all timepoints? |
| --- | --- | --- | --- | --- | --- | --- | --- |
| ITIC-04 | Labrador Retriever | SF | 10 | 38.9 | Left Maxilla/Pulmonary nodule (T2N0M1) | A | Yes |
| ITIC-05 | Burnese Mountain | NM | 7 | 43.4 | Right hindlimb subcutaneous (T2N2M0) | B | Yes |
| ITIC-06 | Min. Poodle mix | NM | 10 | 6.6 | Right Mandible (T2N1M0) | A | No<sup>b</sup> |
| ITIC-07 | Aust. Cattle mix | NM | 13 | 31.2 | Left Maxilla (T2N0M0) | B | Yes |
| ITIC-08 | Beagle mix | NM | 11 | 15.8 | Soft palate (T2N1M0) | B | Yes |
| ITIC-09 | Great Dane | SF | 10 | 51.7 | Left Maxilla (T2N1M1) | A | Yes |
| ITIC-10 | German Shepherd | NM | 9 | 26.3 | Right Maxilla (T2N0M0) | A | Yes |
| ITIC-11 | Mix Breed | NM | 11 | 26.5 | Right Mandible (T2N0M1) | B | Yes |
| ITIC-12 | Pom mix | SF | 11 | 9.0 | Left Maxilla (T3N1M0) | A | No<sup>b</sup> |
| ITIC-13 | Mix Breed | NM | 10 | 7.0 | Cutaneous (TxN1M1) | A | Yes |
| ITIC-14 | Collie mix | SF | 11 | 25.0 | Left Maxilla (T2N1M0) | B | Yes |
| ITIC-15 | Mix Breed | NM | 9 | 36.1 | Lingual (T2N0M0) | B | Yes |

IC, immunocytokine; RT, radiation therapy; SF, spayed female; NM, neutered male
<sup>a</sup>Arm A, a single 8 Gy fraction to the primary tumor followed 5 days later with hu14.18-IL2 IC (12 mg/m<sup>2</sup> for 3 consecutive days); Arm B, three 8 Gy fractions delivered on a

All 12 dogs completed the treatment phase of the protocol, and none experienced significant or unexpected adverse events [38].

### Detection of CAHAs by ELISA

We developed an in-house sandwich ELISA to detect antibodies in canine sera specific to human IgG — the antibody portion of hu14.18 is humanized IgG. For this assay to work, a secondary antibody specific for, or cross-reactive to, canine immunoglobulin is required. Additionally, to avoid the need for multiple secondary reagents, reactivity to mouse immunoglobulin, the species the positive control is raised in, was preferred. We first tested our potential secondary reagents by optimizing their ability to generate a strong signal when detecting mouse anti-human IgG mAb as the positive control (in place of the canine serum), as the mouse anti-human IgG was serving as the positive control reagent, namely the analyte. Our initial “checkerboard” trials tested various concentrations of human IgG coating antigen, mouse anti-human IgG mAb as the analyte, and rabbit anti-dog HRP secondary antibody. As expected, the measured absorbance was dependent on the concentration of the rabbit anti-dog HRP secondary, i.e., absorbance decreased as concentration of the secondary decreased. In contrast, the absorbance was not dependent on the concentration of either the coating human IgG or the mouse anti-human IgG mAb (data not shown). To address these issues, we first troubleshot reagents and/or steps, e.g., testing different blocking buffers, including 5% normal rabbit sera, as well as non-mammalian protein buffers, increasing the blocking time, and testing different wash buffers. These changes did not resolve the high background issue, which was also observed in negative control wells, i.e., in wells incubated with PBS instead of the mouse anti-human IgG mAb.

These data suggested that the rabbit anti-dog HRP secondary may be binding to the human IgG coating on the plate. Immunoglobulins from different species share similar protein structures [42]. Therefore, antibodies against one species are likely to cross-react with a number of other species — a property which, as discussed below, was leveraged to optimize the assay. We adsorbed the rabbit anti-dog secondary antibody with excess human IgG; however, recognition of human IgG by this adsorbed reagent was only slightly minimized (data not shown). We are not aware of a commercially available anti-dog secondary antibody adsorbed to human serum proteins. Using the cross-reactivity of different species’ immunoglobulins, we next tested a commercially available rabbit anti-mouse IgG secondary antibody adsorbed to ensure minimal cross-reactivity with human serum proteins. The human adsorbed rabbit anti-mouse IgG did not bind to plate-bound human IgG while the cross-reactivity to the positive control mouse IgG was maintained. This one reagent then permitted the use of a single secondary antibody to effectively detect both CAHA and mouse-anti-human antibody (MAHA).

Post-treatment sera from seven of twelve dogs tested positive for anti-human IgG antibodies determined via our in-house CAHA ELISA (Fig 2). Pre-treatment sera was used as baseline and as an internal control for each dog’s post-treatment samples. Two of six and five of six dogs in Arm A and Arm B, respectively, demonstrated significant increases in anti-human IgG compared to the pre-treatment absorbance (Fig 2). The post-treatment anti-human IgG was significantly increased compared to pre-treatment anti-human IgG at Day 30 for five dogs and at Day 60 for five dogs. For Arm A, both samples reaching significance were from Day 30 (K906 and K909). The sera from K909 no longer had a significant increase in CAHA at Day 60 and no sera was available for Day 60 CAHA analysis from K906. For Arm B, anti-human IgG antibody levels in three dogs were elevated similarly at Day 30 and Day 60 (K905, K911, K915), one dog had an elevated level at Day 30 that declined by Day 60 (K914), and one had an elevated level that was only detected at Day 60 (K908). No dogs had detectable anti-human IgG antibodies following RT but before IT-IC (Day 1) or at Day 10 post-treatment. There were no substantial differences in anti-human IgG levels between negative control wells with PBS only and pre-treatment samples for the 12 dogs enrolled in this study.

**Fig 2.**
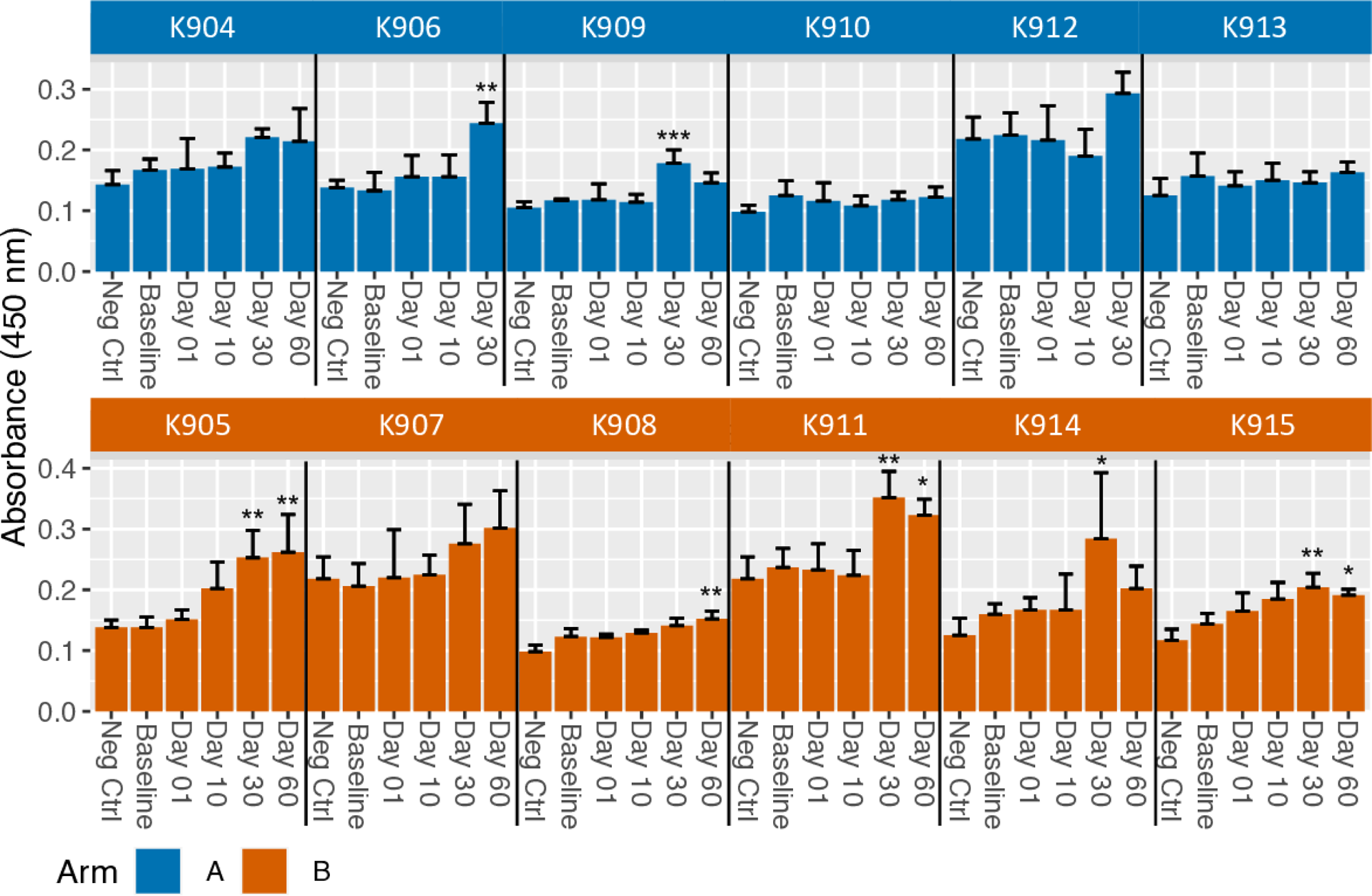
CAHA is induced in some dogs receiving IT-IC injections. Mean CAHA levels, measured as O.D. values by ELISA, from sera assayed in quadruplicate collected at Baseline (before radiation), Day 1 (after radiation, but prior to IT-IC), and Days 10, 30 and 60 (after IT-IC) with standard deviation error bars are shown. Day 60 timepoint was not available for two dogs, K906 and K912, in Arm A. Increases in absorbance from Baseline to Day 1, Day 10, Day 30, or Day 60 for each dog were analyzed and significance indicated as * p < 0.05; ** p < 0.01; *** p < 0.001.

### Sera from dogs treated with hu14.18-IL2 inhibit in vitro binding to GD2

We modified a flow cytometry assay to determine the potential for CAHA in canine sera to inhibit the binding of IC to GD2 on cells [30]. In this assay, sera are first incubated with 14G2a-PE, a mouse mAb that recognizes GD2 at a dilution of ∼1:48.

The mixture is subsequently added to GD2+ M21 cells, and the level of PE fluorescence is determined. 14G2a is an IgG2a-class switch variant of the mouse IgG3 mAb clone 14.18, and thus has the same idiotype as that of the humanized immunocytokine hu14.18-IL2 [46]. Consequently, some of the CAHA in dog sera that recognizes 14G2a may in effect recognize the idiotype shared by 14G2a and hu14.18-IL2, while some of the CAHA might also recognize other regions of the IC. Thus, in this assay, the PE fluorescence of the 14G2a-PE mAb bound to the M21 cells will be decreased if any of the CAHA sin sera from treated subjects recognize the idiotype of 14G2a and effectively block its binding to GD2 on the M21 cells. In contrast, substantial PE fluorescence will be detected if the serum antibodies do not block the binding of 14G2a to GD2.

In preliminary experiments, we assayed healthy canine sera for components in dog serum that might interfere with the flow cytometry assay. Serum is a complex biologic that contains carbohydrates, proteins, and lipids that can interfere with the ability of antibodies to bind their targets. Healthy serum was spiked into PBS and incubated with the 14G2a anti-GD2 mAb before adding the mixture to M21 cells, as described above. Compared to M21 stained in the absence of healthy sera, the addition of sera substantially inhibited 14G2a binding to GD2 in a concentration dependent manner. Substantial inhibition (∼50.2%) was observed at 1:6 ratio of normal sera to PBS whereas at a 1:48 ratio inhibition was largely undetected (> 0.1%) (S1 Fig). As the observed interference of healthy control serum on 14.G2a-PE binding to M21 was ameliorated at the 1:48 ratio, we used this same ratio to test sera from the clinical trial. We included heathy donor sera diluted 1:6 and 1:48 with PBS as positive and negative, respectively, controls in each assay. Each dog’s pre-treatment sample was used as its baseline.

Varying levels of binding inhibition were observed in 10 of 12 dogs (all but K913 and K914) when canine sera were mixed with 14G2a mAb before the mixture was used to stain M21 cells (Table 2, Fig 3, and S2 Fig for select data). The binding was significantly inhibited at Days 10, 30 and 60 compared to pre-treatment in all 10 of these same dogs (save for subjects K906 and K912, that did not have day 60 serum sample collected). GD2 binding was also significantly inhibited by Day 1 sera from 2 of these same 10 dogs, one dog in each Arm, (K910 and K915), but for these 2 dogs, the calculated binding inhibition on Day 1 was not as high as at any of the other days tested for those same 2 dogs. In the remaining 10 dogs, that did not have significant binding inhibition on Day 1, the binding of 14G2a to GD2 was increased at Day 1 for 8 of them (all but K905 and K908), i.e., the % inhibition calculated using the formula in the methods section was negative, corresponding to a slight increase in binding at Day 1 compared to the pre-treatment serum. However, the increase in binding seen on Day 1 (i.e., the negative binding inhibition value) was significant in only one of these eight dogs (K909) (median-8.7%, range-0.3 to-21.2%) (Fig 3 and Table 2). These relatively small positive or negative binding inhibition values detected on Day 1 (prior to any IT-IC treatment, but after RT), which are significant only for 3 dogs (two positive and one negative), were not anticipated and are discussed below; for all 3 of these dogs, all 3 subsequent time points for each dog show higher and statistically significant binding inhibition. In contrast to the 10 dogs that did show significant binding inhibition, sera from the other 2 dogs (K913 and K914), did not inhibit GD2 binding at any timepoint (Fig 3 and Table 2). Select flow cytometry data are shown in S2 Fig.

**Fig 3.**
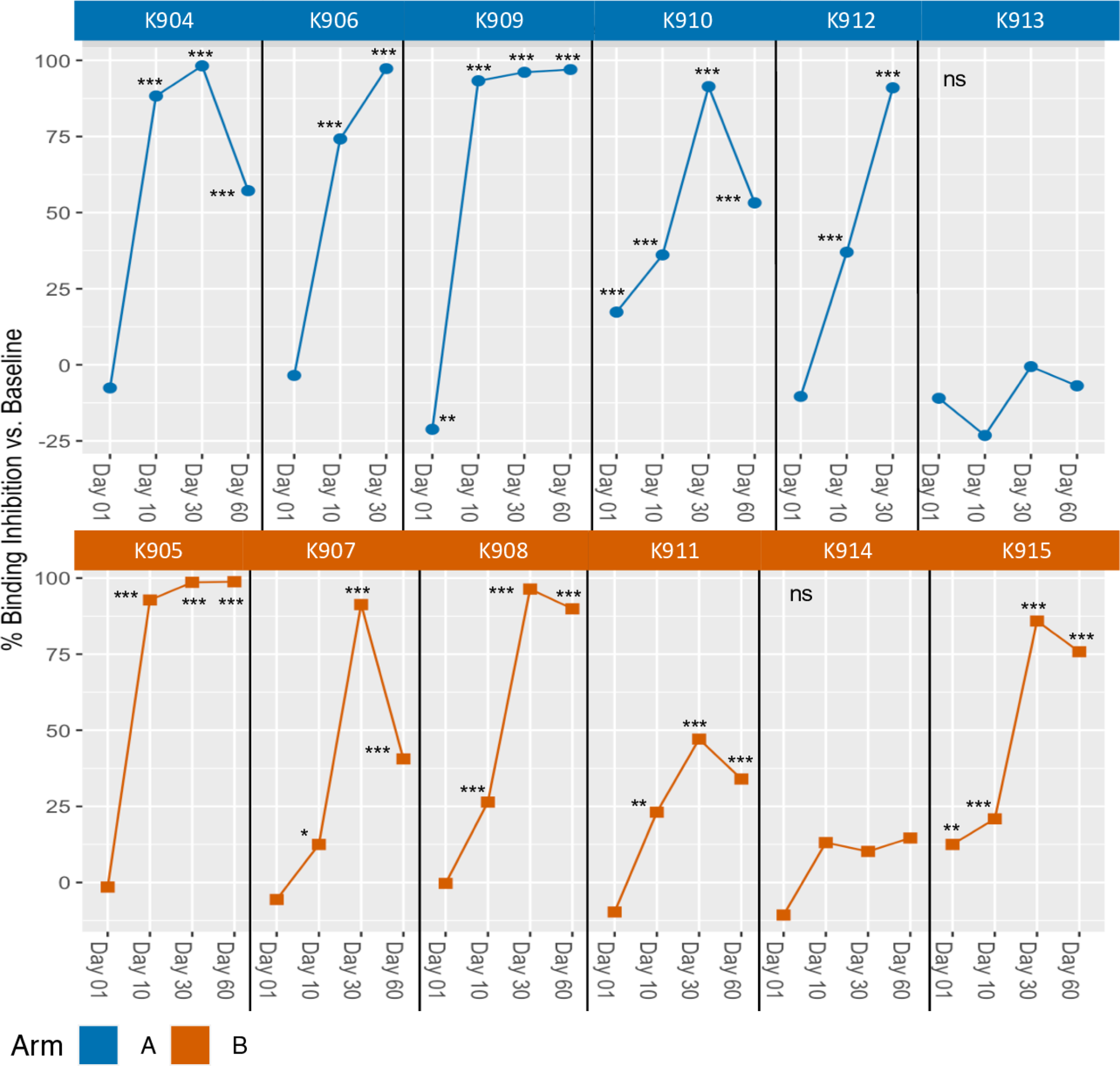
In vitro inhibition of binding of anti-GD2 antibody to GD2 target. The % binding inhibition at various timepoints compared to Baseline. Data shown are calculated from the mean of triplicates as described in Materials and Methods. Sera was collected at Baseline (before radiation), Day 1 (after radiation, but just prior to IT-IC), and Days 10, 30 and 60 (after IT-IC). Timepoints for each dog are presented and connected by a line. Day 60 timepoint was not available for two dogs, K906 and K912, in Arm A. Significance indicated is based on mean of triplicates at Baseline compared to post-treatment timepoints, * p < 0.05; ** p < 0.01; *** p < 0.001.

**Table 2.** Binding Inhibition (%) of anti-GD2 mAb to GD2 target in the presence of canine serum.

| Patient ID | RT Arm | Day 1 | Day 10 | Day 30 | Day 60 |
| --- | --- | --- | --- | --- | --- |
| ITIC-04 | A | -7.6 | 88.3 | 98.2 | 57.2 |
| ITIC-05 | B | -1.5 | 92.8 | 96.6 | 98.8 |
| ITIC-06 | A | -3.5 | 74.2 | 97.3 | n/a |
| ITIC-07 | B | -5.6 | 12.5 | 91.3 | 40.6 |
| ITIC-08 | B | -0.3 | 26.4 | 96.4 | 89.9 |
| ITIC-09 | A | -21.2 | 93.3 | 96.1 | 97.0 |
| ITIC-10 | A | 17.3 | 36.1 | 91.4 | 53.2 |
| ITIC-11 | B | -9.7 | 23.1 | 47.1 | 34.0 |
| ITIC-12 | A | -10.4 | 37.0 | 91.0 | n/a |
| ITIC-13 | A | -11.0 | -23.2 | -0.6 | -6.9 |
| ITIC-14 | B | -10.7 | 13.1 | 10.2 | 14.6 |
| ITIC-15 | B | 12.5 | 20.9 | 85.9 | 75.8 |

mAb, monoclonal antibody; RT, radiation therapy Fluorescently labelled anti-GD2 antibody was incubated with canine serum and then used to stain GD2+ M21 tumor cells. % Binding Inhibition of post-treatment samples compared to baseline samples was determined by flow cytometry as described in Materials and Methods.

### Comparison of CAHA Detection by ELISA and binding inhibition by *in vitro* flow cytometry assay

Significant CAHA at various timepoints was detected by ELISA in 7 of 12 dogs: i) K906 and K909 from Arm A at Day 30; ii) K908 from Arm B at Day 60; iii) K914 from Arm B at Day 30; and iv) K905, K911, and K9 15 from Arm B at Day 30 and Day 60 (Fig 2).

Except for the Day 30 sera from K914, the above sera also significantly inhibited 14G2a from binding to GD2 in the flow cytometry assay (Fig 3). To evaluate the relationship between the ELISA and binding inhibition, the values for each serum sample for each subject are plotted in Fig 4, with the ELISA O.D. value plotted on the Y axis, and the % binding inhibition value plotted on the X-axis. There were no significant correlations between the two assays within or for both arms combined detected. Significant binding inhibition was detected earlier than significant CAHA: i) at Day 1 in 2 dogs; and ii) at Day 10 in 10 dogs. Overall, inhibition was detected in 10 of 12 dogs: i) 5 of 6 Arm A dogs and 5 of 6 Arm B dogs at Day 10; ii) 5 of 6 Arm A dogs and 5 of 6 Arm B dogs at Day 30; iii) 3 of 4 Arm A dogs and 5 of 6 Arm B dogs at Day 60. Sera from the two dogs that did not inhibit binding in the flow cytometry assay (K913 and K914) were also CAHA negative by ELISA except for the Day 30 sera from K914 (Figs 2 and 3). Healthy canine sera at the same dilution as sera from the clinical trial dogs were negative for CAHA and did not inhibit the binding of 14G2a to M21 (Fig B in S1 Fig).

**Fig 4.**
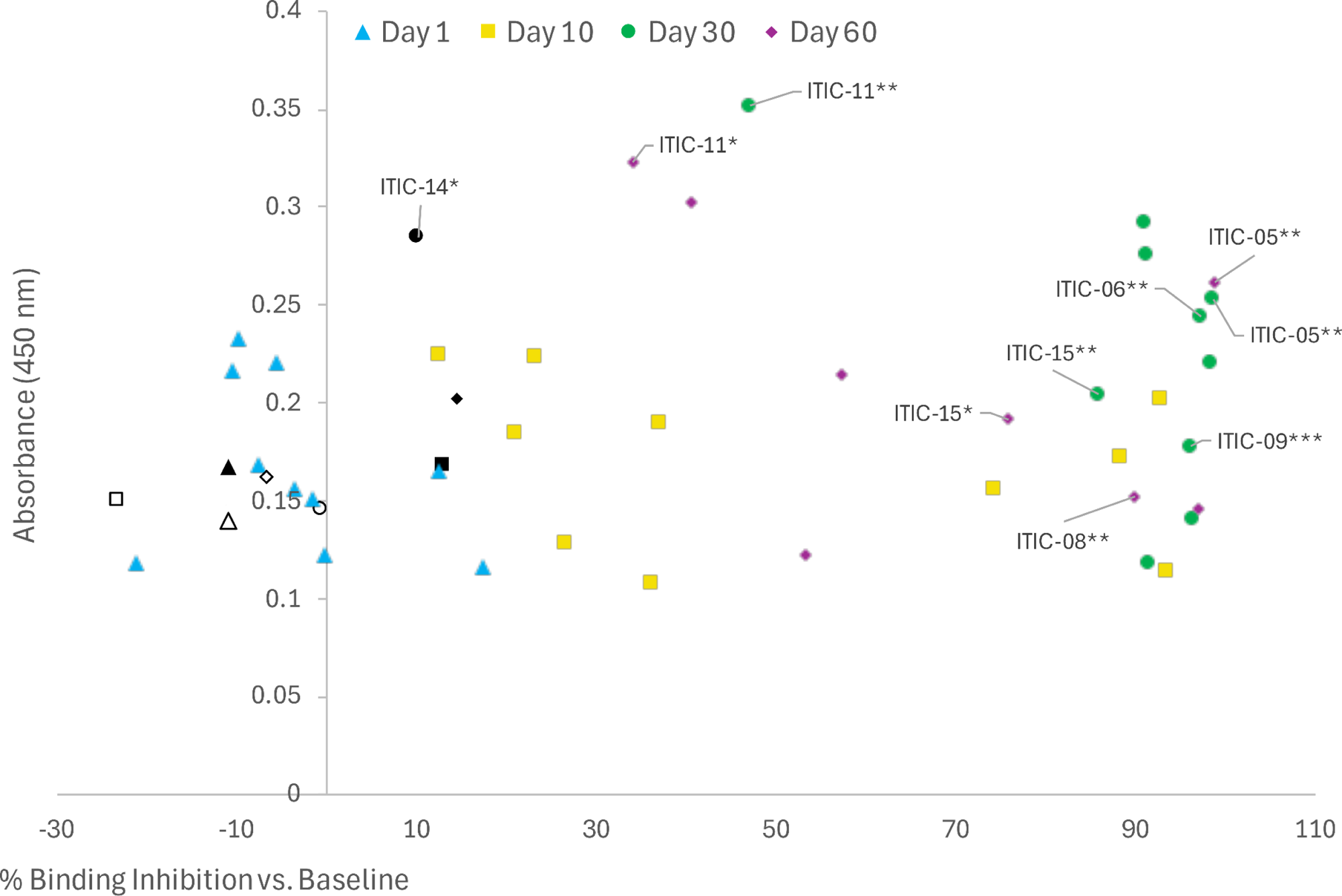
The relationship of CAHA and binding inhibition by *in vitro* flow cytometry assay. The % binding inhibition related to CAHA at various timepoints is shown. Timepoints for each dog are presented. Timepoints with significant CAHA responses are shown with significance indicated, these timepoints also had significant binding inhibition except for K914 Day 30. Sera from 10 of 12 dogs significantly inhibited binding of anti-GD2 antibody to GD2 target at Day 10, 30 and 60 with the exceptions of K913 (shown as open symbols) and K914 (shown as black symbols). Day 1 sera from two dogs (K910 and K915) inhibited binding. A single serum sample, K914 Day 30, showed a significant CAHA response but did not inhibit binding (shown as the black circle).

## Discussion

We developed two distinct assays to monitor and assess the CAHA response in dogs receiving treatment of a readily palpable oral malignant melanoma with local RT plus IT injection of a humanized IC. An ELISA was designed to detect CAHAs specifically targeting human IgG – which is analogous to the Fc portion of hu14.18-IL2 [46]. We also utilized a flow cytometry assay to investigate the inhibition of anti-GD2 mAb binding to GD2 by sera from dogs receiving RT and IT-IC. This approach employed ELISA as a tool to identify the presence of ADA in serum, followed by a subsequent flow cytometry analysis to provide additional characterization of the ADA response.

Although there are commercially available assays to detect various anti-species antibodies, e.g., mouse anti-human (MAHA) and human anti-mouse (HAMA), an off-the-shelf assay is not available to detect CAHA. The absence of a commercially available ELISA specifically designed to detect CAHAs is an unmet need within the comparative immuno-oncology field. This gap becomes increasingly relevant as the canine model gains recognition for its value in studying human disease. The primary objective of developing our in-house CAHA ELISA was to create a monitoring tool that would be easy to use in veterinary and translational research laboratories. Additionally, we sought to develop a cost-effective and flexible assay. The binding specificity of the CAHA could also be explored in more depth by modifying the whole human IgG coating reagent that was used in this report (e.g., whole IgG, Fc fragment, Fab or F(ab’)_2_).

A challenge in developing an ELISA to detect canine antibody that binds human IgG was the lack of commercially available anti-canine IgG HRP conjugates that were cross absorbed against human IgG. This cross-absorption step is crucial for reducing the assay background and ensuring specificity. To address this limitation, we used a rabbit anti-mouse IgG HRP conjugate that had been cross absorbed against human IgG. Our testing during assay optimization demonstrated sufficient cross-reactivity between mouse and canine-derived IgG, enabling us to detect CAHA to human IgG (data not shown). In addition, we included a commercially available mouse IgG specific for human IgG as a control analyte sample for detection by the rabbit anti-mouse HRP in each assay. The selection of antibodies and the incorporation of cross-absorption steps contributed to the assay’s specificity and adaptability for potential use in both veterinary and translational research lab settings.

Our CAHA ELISA is primarily designed to detect free ADAs in serum, by detecting their binding to plate-bound human IgG. In contrast, the in vitro binding inhibition flow cytometry assay allows the ADAs to interact with the soluble mouse 14.G2a mAb (which shares the idiotype of the hu14.18-IL2 IC) in solution, and evaluates detection of GD2 binding in a cellular context, enabling the detection of ADAs that inhibit or interfere with the binding of a therapeutic drug to its specific target in its natural form on the cell surface, e.g., GD2 on tumor cells. This distinction is critical because antigens immobilized to plastic, as in an ELISA, may not accurately mimic the natural conformations of antigens on cell surfaces [47], potentially leading to an underestimation of certain ADAs by this ELISA that flow cytometry can detect.

It is of interest that two dogs (K910 and K915) inhibited the binding of the anti-GD2 mAb to GD2 on target cells at Day 1. The Day 1 sample was collected the day of, but prior to, the first injection of hu14.18-IL2. Therefore, these responses following RT may reflect pre-existing canine anti-human or anti-mouse antibodies, cross species-reactive antibodies, or assay interference [48–50]. The first step in the flow cytometry binding inhibition assay is the incubation of canine sera with the mouse anti-GD2 mAb 14G2a prior to incubating the mixture with the GD2+ M21 cells. Therefore, a possible explanation for the Day 1 binding inhibition is pre-existing canine anti-mouse antibodies as previously reported [49], but these would have needed to appear, or be boosted, following baseline, as the positive binding inhibition is detected in comparison to the baseline value for that same dog. We did not prospectively survey clients regarding their pet’s known allergies or exposures to other mammalian species, e.g., mouse or rabbit, which could potentially induce a humoral response. mAbs of both mouse and rabbit origin are used in our in-house assays and could result in assay background or interference. However, neither of these two dogs were CAHA+ by ELISA at Day 1. Anti-human IgG was not detected by ELISA until Day 30 in any dog, nor from healthy canine sera, suggesting exposure to hu14.18-IL2 as the origin of the antibodies. However, we have not formally tested for pre-existing anti-mouse or anti-rabbit antibodies in these dogs. These seemingly discordant data could reflect the different assay sensitivities and different assay modalities. Moreover, serum from healthy humans and healthy dogs can both contain detectable levels of autoantibodies including rheumatoid factors and anti-idiotypic antibodies which may cause assay interference [48, 50, 51].

Additionally, activation of B cells, generation of anti-tumor antibodies and establishment of B cell memory following a treatment regimen similar to that used in this trial have been reported in a mouse melanoma model [52]. As IgM typically arises early in an immune response, the CAHA response detected at early timepoints (e.g., Day 1, Day 10) in this study may be of IgM isotype which our CAHA ELISA may also detect. in contrast, the flow cytometry assay does not detect a particular antibody isotype and thus may detect ADA despite lack of CAHA detected by ELISA in the same serum. The detection of binding inhibition at early time points may also result from the isotype agnostic nature of the flow cytometry assay. The increase in binding inhibition observed in two dogs (K910 and K915) from baseline to Day 1 was unexpected as the dogs had not been exposed to hu14.18-IL2 at the time of serum collection on either day.

Moreover, the timeframe from baseline to Day 1 in this study was between 5-9 days, representing rapid kinetics for a *de novo* humoral response. However, we have not determined the fine specificity of the antibody responses in this study. All dogs treated in this study received radiation, thus there is potential for radiation-induced antigen shedding by tumor to trigger a humoral response [53]. Tumor-specific and self-specific antibodies have been found in sera of cancer patients in several studies [54, 55]. Thus, the binding inhibition at Day 1 in these two dogs could be due to endogenous cross-reactive antibodies or those recognizing cryptic epitopes or neoantigens [56].

The IC has humanized IL2 linked to the mAb Fc region, thus the IL2 and/or the linker between the Fc and the IL2 may be recognized as foreign in the dog triggering an immune response. None of the dogs in this study experienced significant or unexpected adverse events [38]. However, as dogs received only a single 3-day course of IT-IC, we could not evaluate the impact of induced ADAs on subsequent pharmacokinetics of hu14.18-IL2 or on the potential toxicity during subsequent courses. It will be important to evaluate peak serum levels of hu14.18-IL2 as well as monitor anti-IL2 and anti-linker responses to determine if CAHA+ sera affect antibody-mediated cellular cytotoxicity, as seen in some humans receiving IC treatment [26].

Additional studies could further characterize the binding specificity in terms of distinct IgG regions, e.g., Fc receptor, Fab hinge region [49]. Further, as in humans, IgG is the most prevalent Ig in canine sera, however IgM and IgA are present at lower concentrations [57]. The rabbit anti-mouse IgG HRP secondary reagent used in the ELISA is reactive to both the heavy and light chain of mouse IgG. As such, this reagent may also react with other immunoglobulin classes, e.g., IgM, IgA, since these all share the same light chain. Therefore, subsequent testing could determine the isotype of the CAHA responses, as well as species cross-reactivity – in particular to mouse or rabbit IgG.

The timeline of the anti-human IgG CAHA ELISA response observed in our study (7 of 12 dogs at Day 30) is similar to that observed in a previous study of dogs treated with NHS-IL12, a humanized antibody against necrotic tumor regions linked to humanized IL12. Dogs treated with NHS-IL12 developed ADA against both human IgG (8 of 12 dogs) and against human IL12 (5 of 14 dogs) between 15 to 29 days after treatment [31]. Factors that may explain differences in the ADA timeline between our study and that of Paoloni et al., include the composition and target of the IC, IC dose level, administration route, as well as the assays used to monitor the development of ADA.

## Conclusion

The CAHA detection assays in this report provide tools to monitor parallel canine patient populations receiving treatment with human biopharmaceuticals. These assays should be translatable to other treatment regimens delivering “foreign” protein molecules *in vivo*, as well as to species other than the dog, provided appropriate controls are available. The recent availability of immune checkpoint blockade in the form of a caninized anti-PD1 therapeutic mAb (Gilvetmab, Merck) expands the treatment options for dogs with melanoma. To this end, we are evaluating the addition of immune checkpoint blockade, i.e., anti-PD1, to our novel treatment regimen of RT and IT-IC with hu14.18-IL2 in dogs with melanoma. Local anti-tumor responses as well as systemic immune responses, including the development of CAHA using the assays described herein, will be investigated.

## Acknowledgements

We are grateful for the dogs with melanoma, their caregivers, and referring veterinarians who participated in the clinical trial. We are also thankful for the healthy dogs and their caregivers for donating peripheral blood samples. We thank Rubi Hayim, Emily M. Tomesek, and the UW Veterinary Care Oncology team for caring for the dogs in this study.

## Supporting information

**S1 Fig.**
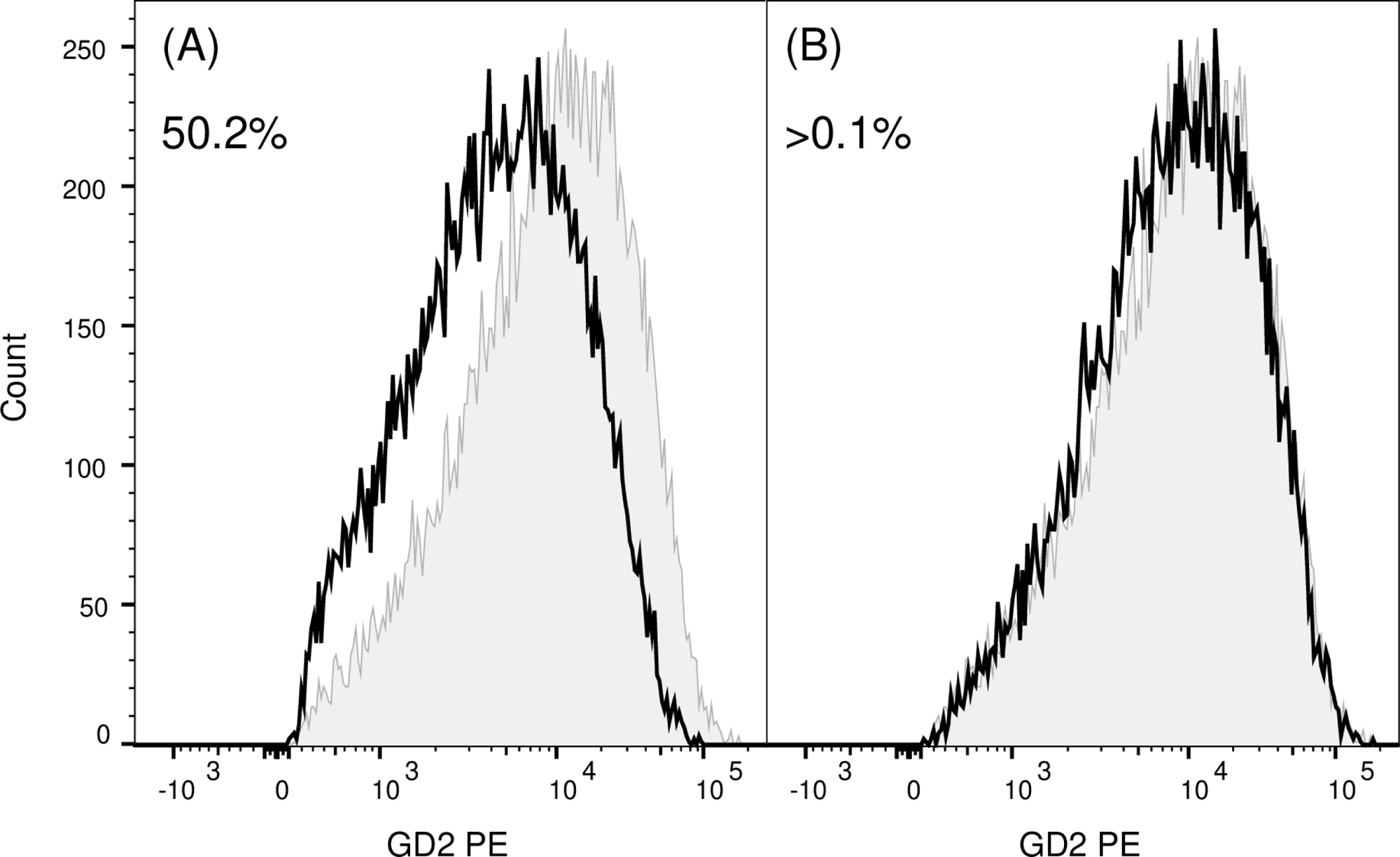
Binding inhibition assay controls. PE-conjugated anti-GD2 antibody 14G2a was mixed with either (A) 20 μl (open histogram with solid line) or (B) 2.5 μl (open histogram with solid line) healthy canine donor sera before staining M21 cells. M21 stained with 14G2a-PE mixed with 20 μl or 2.5 μl PBS as a no-serum control is represented as a gray histogram in both (A) and (B). Data are representative of triplicates. Values represent % binding inhibition compared to no-serum control.

**S2 Fig.**
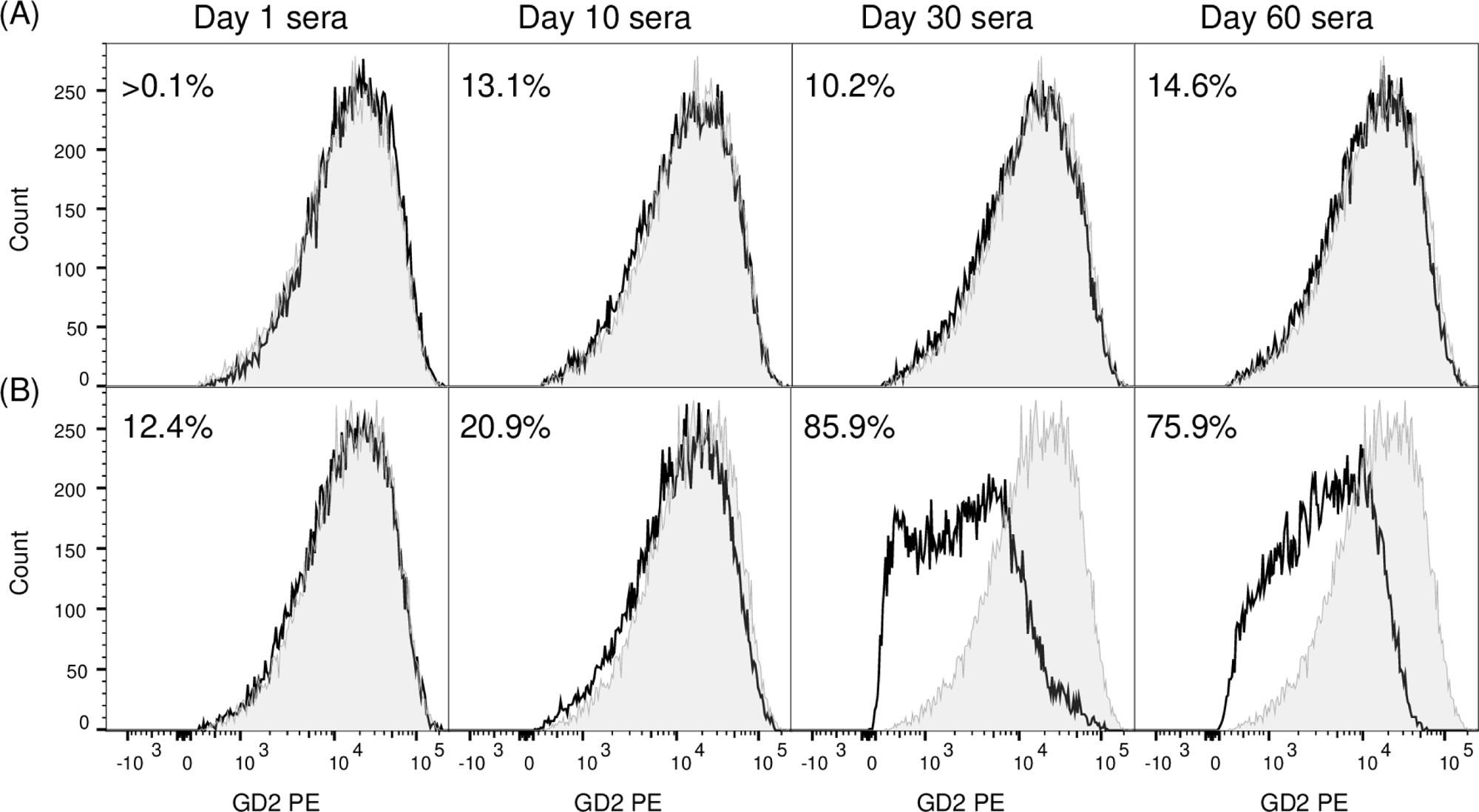
Serum from dogs treated with hu14.18-IL2 shows varied level of inhibition of 14G2a binding to target GD2. Prior to staining M21 cells, PE-conjugated anti-GD2 antibody 14G2a was mixed with baseline/pre-treatment sera or post-treatment sera from different timepoints from (A) K914 or (B) K915. Baseline/pre-treatment data are represented as gray histograms; post-treatment data are represented as open histograms with solid lines. Data are representative of triplicates. Values represent % binding inhibition compared to baseline/pre-treatment control.

## Data availability statement

All relevant data are within the manuscript and its Supporting Information files.

