## Supplementary material for "In vitro detection of canine anti-human antibodies following intratumoral injection of the hu14.18-IL2 immunocytokine in spontaneous canine melanoma": S3_Dataset

| arm | patient.id | tmpt | MFI |
|---|---|---|---|
| A | ITIC-04 | Baseline | 10602 |
| A | ITIC-04 | Baseline | 12184 |
| A | ITIC-04 | Baseline | 13544 |
| A | ITIC-04 | Day 1 | 13327 |
| A | ITIC-04 | Day 1 | 13023 |
| A | ITIC-04 | Day 1 | 12757 |
| A | ITIC-04 | Day 10 | 1145 |
| A | ITIC-04 | Day 10 | 1026 |
| A | ITIC-04 | Day 10 | 2076 |
| A | ITIC-04 | Day 30 | 195 |
| A | ITIC-04 | Day 30 | 191 |
| A | ITIC-04 | Day 30 | 265 |
| A | ITIC-04 | Day 60 | 5393 |
| A | ITIC-04 | Day 60 | 4997 |
| A | ITIC-04 | Day 60 | 5147 |
| B | ITIC-05 | Baseline | 10459 |
| B | ITIC-05 | Baseline | 10578 |
| B | ITIC-05 | Baseline | 11020 |
| B | ITIC-05 | Day 1 | 10110 |
| B | ITIC-05 | Day 1 | 11045 |
| B | ITIC-05 | Day 1 | 11377 |
| B | ITIC-05 | Day 10 | 727 |
| B | ITIC-05 | Day 10 | 722 |
| B | ITIC-05 | Day 10 | 873 |
| B | ITIC-05 | Day 30 | 204 |
| B | ITIC-05 | Day 30 | 121 |
| B | ITIC-05 | Day 30 | 112 |
| B | ITIC-05 | Day 60 | 146 |
| B | ITIC-05 | Day 60 | 118 |
| B | ITIC-05 | Day 60 | 123 |
| A | ITIC-06 | Baseline | 19576 |
| A | ITIC-06 | Baseline | 18941 |
| A | ITIC-06 | Baseline | 18200 |
| A | ITIC-06 | Day 1 | 19995 |
| A | ITIC-06 | Day 1 | 20184 |
| A | ITIC-06 | Day 1 | 18545 |
| A | ITIC-06 | Day 10 | 4701 |
| A | ITIC-06 | Day 10 | 5180 |
| A | ITIC-06 | Day 10 | 4751 |

| | | | |
|---|---|---|---|
| A | ITIC-06 | Day 30 | 516 |
| A | ITIC-06 | Day 30 | 470 |
| A | ITIC-06 | Day 30 | 561 |
| B | ITIC-07 | Baseline | 18633 |
| B | ITIC-07 | Baseline | 17572 |
| B | ITIC-07 | Baseline | 16382 |
| B | ITIC-07 | Day 1 | 17654 |
| B | ITIC-07 | Day 1 | 18808 |
| B | ITIC-07 | Day 1 | 19075 |
| B | ITIC-07 | Day 10 | 14584 |
| B | ITIC-07 | Day 10 | 15746 |
| B | ITIC-07 | Day 10 | 15672 |
| B | ITIC-07 | Day 30 | 1950 |
| B | ITIC-07 | Day 30 | 1237 |
| B | ITIC-07 | Day 30 | 1397 |
| B | ITIC-07 | Day 60 | 11507 |
| B | ITIC-07 | Day 60 | 9534 |
| B | ITIC-07 | Day 60 | 10202 |
| B | ITIC-08 | Baseline | 18676 |
| B | ITIC-08 | Baseline | 18415 |
| B | ITIC-08 | Baseline | 16730 |
| B | ITIC-08 | Day 1 | 16268 |
| B | ITIC-08 | Day 1 | 18115 |
| B | ITIC-08 | Day 1 | 19576 |
| B | ITIC-08 | Day 10 | 13388 |
| B | ITIC-08 | Day 10 | 12815 |
| B | ITIC-08 | Day 10 | 13388 |
| B | ITIC-08 | Day 30 | 614 |
| B | ITIC-08 | Day 30 | 725 |
| B | ITIC-08 | Day 30 | 599 |
| B | ITIC-08 | Day 60 | 2410 |
| B | ITIC-08 | Day 60 | 1680 |
| B | ITIC-08 | Day 60 | 1335 |
| A | ITIC-09 | Baseline | 17246 |
| A | ITIC-09 | Baseline | 18502 |
| A | ITIC-09 | Baseline | 20184 |
| A | ITIC-09 | Day 1 | 21060 |
| A | ITIC-09 | Day 1 | 23538 |
| A | ITIC-09 | Day 1 | 23206 |
| A | ITIC-09 | Day 10 | 1394 |

| | | | |
|---|---|---|---|
| A | ITIC-09 | Day 10 | 1195 |
| A | ITIC-09 | Day 10 | 1136 |
| A | ITIC-09 | Day 30 | 622 |
| A | ITIC-09 | Day 30 | 828 |
| A | ITIC-09 | Day 60 | 487 |
| A | ITIC-09 | Day 60 | 603 |
| A | ITIC-09 | Day 60 | 568 |
| A | ITIC-10 | Baseline | 20666 |
| A | ITIC-10 | Baseline | 21975 |
| A | ITIC-10 | Baseline | 20911 |
| A | ITIC-10 | Day 1 | 18243 |
| A | ITIC-10 | Day 1 | 17006 |
| A | ITIC-10 | Day 1 | 17327 |
| A | ITIC-10 | Day 10 | 14151 |
| A | ITIC-10 | Day 10 | 12296 |
| A | ITIC-10 | Day 10 | 14151 |
| A | ITIC-10 | Day 30 | 1983 |
| A | ITIC-10 | Day 30 | 1950 |
| A | ITIC-10 | Day 30 | 1518 |
| A | ITIC-10 | Day 60 | 10578 |
| A | ITIC-10 | Day 60 | 9773 |
| A | ITIC-10 | Day 60 | 9364 |
| B | ITIC-11 | Baseline | 12073 |
| B | ITIC-11 | Baseline | 14054 |
| B | ITIC-11 | Baseline | 13701 |
| B | ITIC-11 | Day 1 | 13956 |
| B | ITIC-11 | Day 1 | 14857 |
| B | ITIC-11 | Day 1 | 14891 |
| B | ITIC-11 | Day 10 | 9951 |
| B | ITIC-11 | Day 10 | 9074 |
| B | ITIC-11 | Day 10 | 11586 |
| B | ITIC-11 | Day 30 | 7414 |
| B | ITIC-11 | Day 30 | 7284 |
| B | ITIC-11 | Day 30 | 6377 |
| B | ITIC-11 | Day 60 | 7612 |
| B | ITIC-11 | Day 60 | 9664 |
| B | ITIC-11 | Day 60 | 8993 |
| A | ITIC-12 | Baseline | 12727 |
| A | ITIC-12 | Baseline | 13956 |
| A | ITIC-12 | Baseline | 14151 |

| | | | |
|---|---|---|---|
| A | ITIC-12 | Day 1 | 14961 |
| A | ITIC-12 | Day 1 | 14151 |
| A | ITIC-12 | Day 1 | 15967 |
| A | ITIC-12 | Day 10 | 8448 |
| A | ITIC-12 | Day 10 | 9197 |
| A | ITIC-12 | Day 10 | 8081 |
| A | ITIC-12 | Day 30 | 1145 |
| A | ITIC-12 | Day 30 | 1278 |
| A | ITIC-12 | Day 30 | 1246 |
| A | ITIC-13 | Baseline | 12325 |
| A | ITIC-13 | Baseline | 13956 |
| A | ITIC-13 | Baseline | 13357 |
| A | ITIC-13 | Day 1 | 14754 |
| A | ITIC-13 | Day 1 | 13419 |
| A | ITIC-13 | Day 1 | 15819 |
| A | ITIC-13 | Day 10 | 15856 |
| A | ITIC-13 | Day 10 | 17531 |
| A | ITIC-13 | Day 10 | 15455 |
| A | ITIC-13 | Day 30 | 10723 |
| A | ITIC-13 | Day 30 | 14054 |
| A | ITIC-13 | Day 30 | 15100 |
| A | ITIC-13 | Day 60 | 13204 |
| A | ITIC-13 | Day 60 | 15030 |
| A | ITIC-13 | Day 60 | 14151 |
| B | ITIC-14 | Baseline | 15312 |
| B | ITIC-14 | Baseline | 14857 |
| B | ITIC-14 | Baseline | 16887 |
| B | ITIC-14 | Day 1 | 16966 |
| B | ITIC-14 | Day 1 | 18720 |
| B | ITIC-14 | Day 1 | 16420 |
| B | ITIC-14 | Day 10 | 13796 |
| B | ITIC-14 | Day 10 | 13481 |
| B | ITIC-14 | Day 10 | 13606 |
| B | ITIC-14 | Day 30 | 13388 |
| B | ITIC-14 | Day 30 | 14483 |
| B | ITIC-14 | Day 30 | 14382 |
| B | ITIC-14 | Day 60 | 13481 |
| B | ITIC-14 | Day 60 | 11693 |
| B | ITIC-14 | Day 60 | 14995 |
| B | ITIC-15 | Baseline | 16004 |

| | | | |
|---|---|---|---|
| B | ITIC-15 | Baseline | 17696 |
| B | ITIC-15 | Baseline | 16497 |
| B | ITIC-15 | Day 1 | 15065 |
| B | ITIC-15 | Day 1 | 14021 |
| B | ITIC-15 | Day 1 | 14857 |
| B | ITIC-15 | Day 10 | 12496 |
| B | ITIC-15 | Day 10 | 13512 |
| B | ITIC-15 | Day 10 | 13701 |
| B | ITIC-15 | Day 30 | 3070 |
| B | ITIC-15 | Day 30 | 2095 |
| B | ITIC-15 | Day 30 | 1931 |
| B | ITIC-15 | Day 60 | 4116 |
| B | ITIC-15 | Day 60 | 4246 |
| B | ITIC-15 | Day 60 | 3761 |

| arm | patient.id | tmpt | OD |
|---|---|---|---|
| A | ITIC-04 | Neg Control | 0.169 |
| A | ITIC-04 | Neg Control | 0.123 |
| A | ITIC-04 | Neg Control | 0.125 |
| A | ITIC-04 | Neg Control | 0.155 |
| A | ITIC-04 | Baseline | 0.174 |
| A | ITIC-04 | Baseline | 0.183 |
| A | ITIC-04 | Baseline | 0.171 |
| A | ITIC-04 | Baseline | 0.142 |
| A | ITIC-04 | Day 1 | 0.185 |
| A | ITIC-04 | Day 1 | 0.174 |
| A | ITIC-04 | Day 1 | 0.099 |
| A | ITIC-04 | Day 1 | 0.217 |
| A | ITIC-04 | Day 10 | 0.191 |
| A | ITIC-04 | Day 10 | 0.162 |
| A | ITIC-04 | Day 10 | 0.191 |
| A | ITIC-04 | Day 10 | 0.145 |
| A | ITIC-04 | Day 30 | 0.213 |
| A | ITIC-04 | Day 30 | 0.206 |
| A | ITIC-04 | Day 30 | 0.226 |
| A | ITIC-04 | Day 30 | 0.238 |
| A | ITIC-04 | Day 60 | 0.287 |
| A | ITIC-04 | Day 60 | 0.222 |
| A | ITIC-04 | Day 60 | 0.163 |
| A | ITIC-04 | Day 60 | 0.186 |
| B | ITIC-05 | Neg Control | 0.148 |
| B | ITIC-05 | Neg Control | 0.148 |
| B | ITIC-05 | Neg Control | 0.132 |
| B | ITIC-05 | Neg Control | 0.124 |
| B | ITIC-05 | Baseline | 0.116 |
| B | ITIC-05 | Baseline | 0.137 |
| B | ITIC-05 | Baseline | 0.153 |
| B | ITIC-05 | Baseline | 0.148 |
| B | ITIC-05 | Day 1 | 0.135 |
| B | ITIC-05 | Day 1 | 0.155 |
| B | ITIC-05 | Day 1 | 0.142 |
| B | ITIC-05 | Day 1 | 0.172 |
| B | ITIC-05 | Day 10 | 0.167 |
| B | ITIC-05 | Day 10 | 0.233 |
| B | ITIC-05 | Day 10 | 0.248 |

| | | | |
|---|---|---|---|
| B | ITIC-05 | Day 10 | 0.162 |
| B | ITIC-05 | Day 30 | 0.250 |
| B | ITIC-05 | Day 30 | 0.194 |
| B | ITIC-05 | Day 30 | 0.300 |
| B | ITIC-05 | Day 30 | 0.268 |
| B | ITIC-05 | Day 60 | 0.332 |
| B | ITIC-05 | Day 60 | 0.199 |
| B | ITIC-05 | Day 60 | 0.221 |
| B | ITIC-05 | Day 60 | 0.295 |
| A | ITIC-06 | Neg Control | 0.148 |
| A | ITIC-06 | Neg Control | 0.148 |
| A | ITIC-06 | Neg Control | 0.132 |
| A | ITIC-06 | Neg Control | 0.124 |
| A | ITIC-06 | Baseline | 0.176 |
| A | ITIC-06 | Baseline | 0.127 |
| A | ITIC-06 | Baseline | 0.112 |
| A | ITIC-06 | Baseline | 0.115 |
| A | ITIC-06 | Day 1 | 0.112 |
| A | ITIC-06 | Day 1 | 0.176 |
| A | ITIC-06 | Day 1 | 0.191 |
| A | ITIC-06 | Day 1 | 0.145 |
| A | ITIC-06 | Day 10 | 0.148 |
| A | ITIC-06 | Day 10 | 0.153 |
| A | ITIC-06 | Day 10 | 0.206 |
| A | ITIC-06 | Day 10 | 0.119 |
| A | ITIC-06 | Day 30 | 0.253 |
| A | ITIC-06 | Day 30 | 0.286 |
| A | ITIC-06 | Day 30 | 0.230 |
| A | ITIC-06 | Day 30 | 0.208 |
| B | ITIC-07 | Neg Control | 0.237 |
| B | ITIC-07 | Neg Control | 0.243 |
| B | ITIC-07 | Neg Control | 0.165 |
| B | ITIC-07 | Neg Control | 0.228 |
| B | ITIC-07 | Baseline | 0.254 |
| B | ITIC-07 | Baseline | 0.165 |
| B | ITIC-07 | Baseline | 0.208 |
| B | ITIC-07 | Baseline | 0.197 |
| B | ITIC-07 | Day 1 | 0.318 |
| B | ITIC-07 | Day 1 | 0.244 |
| B | ITIC-07 | Day 1 | 0.181 |

| | | | |
|---|---|---|---|
| B | ITIC-07 | Day 1 | 0.138 |
| B | ITIC-07 | Day 10 | 0.261 |
| B | ITIC-07 | Day 10 | 0.229 |
| B | ITIC-07 | Day 10 | 0.224 |
| B | ITIC-07 | Day 10 | 0.184 |
| B | ITIC-07 | Day 30 | 0.213 |
| B | ITIC-07 | Day 30 | 0.254 |
| B | ITIC-07 | Day 30 | 0.272 |
| B | ITIC-07 | Day 30 | 0.366 |
| B | ITIC-07 | Day 60 | 0.275 |
| B | ITIC-07 | Day 60 | 0.393 |
| B | ITIC-07 | Day 60 | 0.274 |
| B | ITIC-07 | Day 60 | 0.266 |
| B | ITIC-08 | Neg Control | 0.105 |
| B | ITIC-08 | Neg Control | 0.089 |
| B | ITIC-08 | Neg Control | 0.087 |
| B | ITIC-08 | Neg Control | 0.110 |
| B | ITIC-08 | Baseline | 0.113 |
| B | ITIC-08 | Baseline | 0.139 |
| B | ITIC-08 | Baseline | 0.126 |
| B | ITIC-08 | Baseline | 0.113 |
| B | ITIC-08 | Day 1 | 0.120 |
| B | ITIC-08 | Day 1 | 0.126 |
| B | ITIC-08 | Day 1 | 0.127 |
| B | ITIC-08 | Day 1 | 0.116 |
| B | ITIC-08 | Day 10 | 0.122 |
| B | ITIC-08 | Day 10 | 0.130 |
| B | ITIC-08 | Day 10 | 0.134 |
| B | ITIC-08 | Day 10 | 0.130 |
| B | ITIC-08 | Day 30 | 0.124 |
| B | ITIC-08 | Day 30 | 0.150 |
| B | ITIC-08 | Day 30 | 0.147 |
| B | ITIC-08 | Day 30 | 0.144 |
| B | ITIC-08 | Day 60 | 0.166 |
| B | ITIC-08 | Day 60 | 0.160 |
| B | ITIC-08 | Day 60 | 0.142 |
| B | ITIC-08 | Day 60 | 0.141 |
| A | ITIC-09 | Neg Control | 0.105 |
| A | ITIC-09 | Neg Control | 0.099 |
| A | ITIC-09 | Neg Control | 0.097 |

| | | | |
|---|---|---|---|
| A | ITIC-09 | Neg Control | 0.120 |
| A | ITIC-09 | Baseline | 0.115 |
| A | ITIC-09 | Baseline | 0.118 |
| A | ITIC-09 | Baseline | 0.114 |
| A | ITIC-09 | Baseline | 0.119 |
| A | ITIC-09 | Day 1 | 0.153 |
| A | ITIC-09 | Day 1 | 0.094 |
| A | ITIC-09 | Day 1 | 0.106 |
| A | ITIC-09 | Day 1 | 0.118 |
| A | ITIC-09 | Day 10 | 0.129 |
| A | ITIC-09 | Day 10 | 0.118 |
| A | ITIC-09 | Day 10 | 0.096 |
| A | ITIC-09 | Day 10 | 0.114 |
| A | ITIC-09 | Day 30 | 0.181 |
| A | ITIC-09 | Day 30 | 0.208 |
| A | ITIC-09 | Day 30 | 0.158 |
| A | ITIC-09 | Day 30 | 0.164 |
| A | ITIC-09 | Day 60 | 0.157 |
| A | ITIC-09 | Day 60 | 0.162 |
| A | ITIC-09 | Day 60 | 0.127 |
| A | ITIC-09 | Day 60 | 0.139 |
| A | ITIC-10 | Neg Control | 0.105 |
| A | ITIC-10 | Neg Control | 0.089 |
| A | ITIC-10 | Neg Control | 0.087 |
| A | ITIC-10 | Neg Control | 0.110 |
| A | ITIC-10 | Baseline | 0.139 |
| A | ITIC-10 | Baseline | 0.130 |
| A | ITIC-10 | Baseline | 0.089 |
| A | ITIC-10 | Baseline | 0.140 |
| A | ITIC-10 | Day 1 | 0.114 |
| A | ITIC-10 | Day 1 | 0.159 |
| A | ITIC-10 | Day 1 | 0.091 |
| A | ITIC-10 | Day 1 | 0.101 |
| A | ITIC-10 | Day 10 | 0.126 |
| A | ITIC-10 | Day 10 | 0.116 |
| A | ITIC-10 | Day 10 | 0.098 |
| A | ITIC-10 | Day 10 | 0.092 |
| A | ITIC-10 | Day 30 | 0.128 |
| A | ITIC-10 | Day 30 | 0.103 |
| A | ITIC-10 | Day 30 | 0.124 |

| | | | |
|---|---|---|---|
| A | ITIC-10 | Day 60 | 0.139 |
| A | ITIC-10 | Day 60 | 0.122 |
| A | ITIC-10 | Day 60 | 0.106 |
| B | ITIC-11 | Neg Control | 0.237 |
| B | ITIC-11 | Neg Control | 0.243 |
| B | ITIC-11 | Neg Control | 0.165 |
| B | ITIC-11 | Neg Control | 0.228 |
| B | ITIC-11 | Baseline | 0.218 |
| B | ITIC-11 | Baseline | 0.270 |
| B | ITIC-11 | Baseline | 0.257 |
| B | ITIC-11 | Baseline | 0.204 |
| B | ITIC-11 | Day 1 | 0.179 |
| B | ITIC-11 | Day 1 | 0.261 |
| B | ITIC-11 | Day 1 | 0.274 |
| B | ITIC-11 | Day 1 | 0.217 |
| B | ITIC-11 | Day 10 | 0.242 |
| B | ITIC-11 | Day 10 | 0.250 |
| B | ITIC-11 | Day 10 | 0.240 |
| B | ITIC-11 | Day 10 | 0.162 |
| B | ITIC-11 | Day 30 | 0.338 |
| B | ITIC-11 | Day 30 | 0.413 |
| B | ITIC-11 | Day 30 | 0.314 |
| B | ITIC-11 | Day 30 | 0.342 |
| B | ITIC-11 | Day 60 | 0.292 |
| B | ITIC-11 | Day 60 | 0.354 |
| B | ITIC-11 | Day 60 | 0.329 |
| B | ITIC-11 | Day 60 | 0.316 |
| A | ITIC-12 | Neg Control | 0.237 |
| A | ITIC-12 | Neg Control | 0.243 |
| A | ITIC-12 | Neg Control | 0.165 |
| A | ITIC-12 | Neg Control | 0.228 |
| A | ITIC-12 | Baseline | 0.257 |
| A | ITIC-12 | Baseline | 0.234 |
| A | ITIC-12 | Baseline | 0.234 |
| A | ITIC-12 | Baseline | 0.172 |
| A | ITIC-12 | Day 1 | 0.272 |
| A | ITIC-12 | Day 1 | 0.259 |
| A | ITIC-12 | Day 1 | 0.157 |
| A | ITIC-12 | Day 1 | 0.177 |
| A | ITIC-12 | Day 10 | 0.246 |

| | | | |
|---|---|---|---|
| A | ITIC-12 | Day 10 | 0.159 |
| A | ITIC-12 | Day 10 | 0.151 |
| A | ITIC-12 | Day 10 | 0.204 |
| A | ITIC-12 | Day 30 | 0.258 |
| A | ITIC-12 | Day 30 | 0.323 |
| A | ITIC-12 | Day 30 | 0.267 |
| A | ITIC-12 | Day 30 | 0.323 |
| A | ITIC-13 | Neg Control | 0.159 |
| A | ITIC-13 | Neg Control | 0.129 |
| A | ITIC-13 | Neg Control | 0.123 |
| A | ITIC-13 | Neg Control | 0.090 |
| A | ITIC-13 | Baseline | 0.113 |
| A | ITIC-13 | Baseline | 0.183 |
| A | ITIC-13 | Baseline | 0.138 |
| A | ITIC-13 | Baseline | 0.194 |
| A | ITIC-13 | Day 1 | 0.122 |
| A | ITIC-13 | Day 1 | 0.174 |
| A | ITIC-13 | Day 1 | 0.131 |
| A | ITIC-13 | Day 1 | 0.134 |
| A | ITIC-13 | Day 10 | 0.133 |
| A | ITIC-13 | Day 10 | 0.135 |
| A | ITIC-13 | Day 10 | 0.141 |
| A | ITIC-13 | Day 10 | 0.192 |
| A | ITIC-13 | Day 30 | 0.139 |
| A | ITIC-13 | Day 30 | 0.129 |
| A | ITIC-13 | Day 30 | 0.171 |
| A | ITIC-13 | Day 30 | 0.144 |
| A | ITIC-13 | Day 60 | 0.141 |
| A | ITIC-13 | Day 60 | 0.172 |
| A | ITIC-13 | Day 60 | 0.156 |
| A | ITIC-13 | Day 60 | 0.181 |
| B | ITIC-14 | Neg Control | 0.159 |
| B | ITIC-14 | Neg Control | 0.129 |
| B | ITIC-14 | Neg Control | 0.123 |
| B | ITIC-14 | Neg Control | 0.090 |
| B | ITIC-14 | Baseline | 0.164 |
| B | ITIC-14 | Baseline | 0.134 |
| B | ITIC-14 | Baseline | 0.175 |
| B | ITIC-14 | Baseline | 0.165 |
| B | ITIC-14 | Day 1 | 0.169 |

| | | | |
|---|---|---|---|
| B | ITIC-14 | Day 1 | 0.166 |
| B | ITIC-14 | Day 1 | 0.191 |
| B | ITIC-14 | Day 1 | 0.143 |
| B | ITIC-14 | Day 10 | 0.106 |
| B | ITIC-14 | Day 10 | 0.128 |
| B | ITIC-14 | Day 10 | 0.208 |
| B | ITIC-14 | Day 10 | 0.228 |
| B | ITIC-14 | Day 30 | 0.444 |
| B | ITIC-14 | Day 30 | 0.198 |
| B | ITIC-14 | Day 30 | 0.258 |
| B | ITIC-14 | Day 30 | 0.238 |
| B | ITIC-14 | Day 60 | 0.152 |
| B | ITIC-14 | Day 60 | 0.225 |
| B | ITIC-14 | Day 60 | 0.195 |
| B | ITIC-14 | Day 60 | 0.235 |
| B | ITIC-15 | Neg Control | 0.129 |
| B | ITIC-15 | Neg Control | 0.123 |
| B | ITIC-15 | Neg Control | 0.090 |
| B | ITIC-15 | Neg Control | 0.125 |
| B | ITIC-15 | Baseline | 0.154 |
| B | ITIC-15 | Baseline | 0.164 |
| B | ITIC-15 | Baseline | 0.128 |
| B | ITIC-15 | Baseline | 0.132 |
| B | ITIC-15 | Day 1 | 0.168 |
| B | ITIC-15 | Day 1 | 0.167 |
| B | ITIC-15 | Day 1 | 0.199 |
| B | ITIC-15 | Day 1 | 0.126 |
| B | ITIC-15 | Day 10 | 0.211 |
| B | ITIC-15 | Day 10 | 0.204 |
| B | ITIC-15 | Day 10 | 0.169 |
| B | ITIC-15 | Day 10 | 0.155 |
| B | ITIC-15 | Day 30 | 0.189 |
| B | ITIC-15 | Day 30 | 0.206 |
| B | ITIC-15 | Day 30 | 0.237 |
| B | ITIC-15 | Day 30 | 0.186 |
| B | ITIC-15 | Day 60 | 0.201 |
| B | ITIC-15 | Day 60 | 0.199 |
| B | ITIC-15 | Day 60 | 0.180 |
| B | ITIC-15 | Day 60 | 0.187 |
